# The *FLOWERING LOCUS T LIKE 2-1* gene of *Chenopodium* triggers precocious flowering in Arabidopsis seedlings

**DOI:** 10.1101/2022.12.20.521231

**Authors:** Oushadee A. J. Abeyawardana, Tomáš Moravec, Manuela Krüger, Claudia Belz, David Gutierrez-Larruscain, Zuzana Vondráková, Kateřina Eliášová, Helena Štorchová

## Abstract

The *FLOWERING LOCUS T* (*FT)* gene is the essential integrator of flowering regulatory pathways in angiosperms. The paralogs of the *FT* gene may perform antagonistic functions, as exemplified by *BvFT1*, that suppresses flowering in *Beta vulgaris*, unlike the paralogous activator *BvFT2*. The roles of *FT* genes in other amaranths were less investigated. Here, we transformed *Arabidopsis thaliana* with the *FLOWERING LOCUS T like* (*FTL*) genes of *Chenopodium* and found, that both *FTL1* and *FTL2-1* accelerated flowering, despite having been the homologs of the *Beta vulgaris* floral promoter and suppressor, respectively. The floral promotive effect of *FTL2-1* was so strong that it caused lethality when overexpressed under the *35S* promoter. *FTL2-1* placed in inducible cassette accelerated flowering after the induction with methoxyphenozide. The occasional expression of *FTL2-1* led to precocious flowering in some primary transformants even without chemical induction. After the *FTL* gene duplication in Amaranthaceae, the *FTL1* copy maintained the role of floral activator. The second copy *FTL2* underwent subsequent duplication and functional diversification, which enabled to control the onset of flowering in amaranths to adapt to variable environments.

**HIGHLIGHT:** The *FLOWERING LOCUS T like 2-1* (*FTL 2-1*) gene of *Chenopodium* acts as a strong activator of flowering in Arabidopsis, despite being a homolog of floral repressor *BvFT1*.

## INTRODUCTION

The decision when to flower is one of the most important commitments in a plant’s life since it directly impacts evolutionary success of the species. The formation of flowers and seeds requires re-allocation of resources from the entire plant to maximize reproductive success, which is often followed by senescence in annuals. The proper timing of flowering helps the plant to balance reproductive cost and benefit. The onset of flowering is precisely controlled by environmental conditions (daylength, cold temperature in winter, ambient temperature, abiotic stress) as well as on endogenous factors (age, phytohormone concentrations, carbohydrate status) (Andres and Coupland, 2012; Riboni *et al*., 2013; Hyun *et al*., 2016).

The central position at the crossover of the signalling pathways is occupied by the FLOWERING LOCUS T (FT) protein, which is the important part of the long-sought florigen (Chailakhyan, 1936) in *Arabidopsis thaliana* (hereafter Arabidopsis) (Corbesier *et al*., 2007; Jaeger and Wigge 2007) and other species (Tamaki *et al*., 2007; Hayama *et al*., 2007). The FT protein is produced in the phloem companion cells of the leaves and transported to the apical meristem to trigger flowering (Mathieu *et al*., 2007). The *FT* gene underwent duplications during the evolution of angiosperms and its paralogous copies occasionally acquired the opposite function as flowering suppressors. The pair of floral integrators in *Beta vulgaris*, the sugar beet, includes the BvFT2 protein as a floral promoter and BvFT1 as a floral repressor (Pin *et al*., 2010), exemplifies this dual functionality of FT. The *BvFT1* and *BvFT2* genes repressed or promoted flowering, respectively, when ectopically expressed in sugar beet and Arabidopsis. The reversal from the activation to the inhibition of flowering was caused by three amino acid substitutions in the functional domain of the fourth exon of the *BvFT* genes (Pin *et al*., 2010).

The orthologs of *BvFT2* and *BvFT1* were found in all members of the family Amaranthaceae so far analyzed (Drabešová *et al*., 2016). The *CrFTL1* gene in *Oxybasis rubra* (syn. *Chenopodium rubrum;* Cháb *et al*., 2008) promoted flowering in Arabidopsis in the same way as its sugar beet ortholog *BvFT2* (Drabešová *et al*., 2014). After the early duplication, which gave rise to the *FT1* and the *FT2* paralogs, a subsequent gene duplication took place after the ancestor of *Beta* had diverged from the ancestors of *Oxybasis* and *Chenopodium*. This event generated two *FTL2* copies, *FTL2-1* and *FTL2-2*, which are next to each other in the quinoa (*Chenopodium quinoa*) genome (Jarvis *et al*., 2017; Štorchová, 2021) as evidence of this tandem duplication.

Unlike the *BvFT1* gene, which was shown to act as floral repressor (Pin *et al*., 2010), the function of its homologs in *Chenopodium* is little known. The expression of the *FTL* genes in the course of floral induction was investigated in *C. ficifolium* and *C. suecicum* (Štorchová *et al*., 2019), close diploid relatives of the important crop quinoa (Štorchová *et al*., 2015; Walsh *et al*., 2015). Whereas *CsFTL2-1* in *C. suecicum* was highly activated by short days, inducing flowering, negligible expression of this gene was observed in *C. ficifolium* under both short and long photoperiods. The low expression of *CfFTL2-1* was particularly noteworthy in the long-day accession *C. ficifolium* 283, which flowered earlier under long days without the apparent activation of any *CfFTL* gene (Štorchová *et al*., 2019). The *CrFTL2-1* homolog in *O. rubra* was completely silenced (Štorchová *et al*., 2019), which excludes any role in floral induction in this species. Thus, the expression of the *FTL2-1* paralog varied among the *Chenopodium-Oxybasis* species and accessions.

The second paralog *FTL2-2* varied in expression profiles too. It was strongly upregulated in *C. suecicum* and in the long-day accession *C. ficifolium* 283 under the floral induction conditions (Štorchová *et al*., 2019). In contrast, the *CrFTL2-2* gene of *O. rubra* exhibited invariant expression, not correlated with flowering. It also did not promote flowering in Arabidopsis, which indicated no participation in floral transition (Drabešová *et al*., 2014). The *FTL2-2* gene underwent dynamic structural evolution. Unlike the *FTL2-1* paralog, which contains four conserved exons and three introns similarly to the other angiosperm *FT* genes, the *FTL2-2* acquired an additional exon and intron (Drabešová *et al*., 2016). Whereas the complete *FTL2-2* gene exists in *O. rubra*. and *C. suecicum*, the large deletion of 130 bp shortened the fourth exon of *CfFTL2-2* in *C. ficifolium* 283 and the entire *CfFTL2-2* gene was deleted in *C. ficifolium* 459. The changes in gene expression and structure, which affected *FTL2* paralogs after the duplication, might have influenced their function and led to sub-or neo-functionalization.

To better understand the function of the *Chenopodium FTL* genes, we transferred the *CfFTL1, CfFTL2-1*, and *CfFTL2-2* genes of *C. ficifolium* to both wild types and *ft*^*-*^ mutants of Arabidopsis and analyzed the flowering phenotypes of the transformants. The *CfFTL1* overexpression accelerated flowering in all Arabidopsis recipients, while the *CfFTL2-2* overexpression had no effect on flowering. Surprisingly, *CfFTL2-1* overexpression was lethal in Arabidopsis and the vector with inducible *CfFTL2-1* had to be constructed to observe the impact of this gene on flowering in Arabidopsis after chemical induction. Our results indicate that *CfFTL1* and *CfFTL2-1* play positive roles in floral promotion.

## MATERIALS AND METHODS

### Preparation of gene constructs for the transformation of Arabidopsis

All constructs used in this work were assembled using the GoldenBraid standard (Sarrion-Perdigones *et al*., 2011). The sequences of the *Chenopodium FTL* genes can be found under the following GenBank accession numbers: *CfFTL1* - MK212025; *CfFTL2*-*1* - MK212027; *CfFTL2*-*2* - MK212026, *CqFTL2-1* - XM_021919867). The open reading frames (ORF) were amplified from *C. ficifolium* cDNA using Phusion polymerase (Thermo Scientific) and primers designed using the GB-domesticator on the GBcloning website (https://gbcloning.upv.es) (**Supplementary Table S1**). Forty ng of the amplified and column-purified (Qiagen) DNA was cloned into the universal domestication plasmid pUPD2 by restriction ligation reaction with BsmBI and T4 ligase (both Thermo Scientific), and selected clones were verified by Sanger sequencing (Eurofins, Germany). The first set of plasmids was designated for constitutive expression of the respective *CfFTL* gene in Arabidopsis. In these vectors the *CfFTL* ORFs were under transcriptional control of the *CaMV 35S* promoter and terminator. The expression levels were increased by Tobacco mosaic virus Omega leader sequence. For inducible expression we modified the methoxyfenozide inducible system VGE (Semenyuk *et al*., 2010) to comply with GoldenBraid standard. The cassette containing the inducible CfFTL2-1 gene was flanked by two tobacco Matrix attachment region (MAR) elements TM2 (Zhang *et al*., 2002) and RB7 (Allen *et al*., 1996). They were designed to reduce position effect and stabilize the variation of transgene expression among individual transgenic lines. They also reduced the likelihood that the transgene might trigger gene silencing, resulting in a gradual loss of transgene expression in T2 and further generations (**Fig. 1**). All used components are summarized in **Supplementary Table S2**. The final binary constructs used for Arabidopsis transformation were assembled using the extended set of vectors alpha 11 - 14 (Dušek *et al*., 2020).

**Figure 1.**
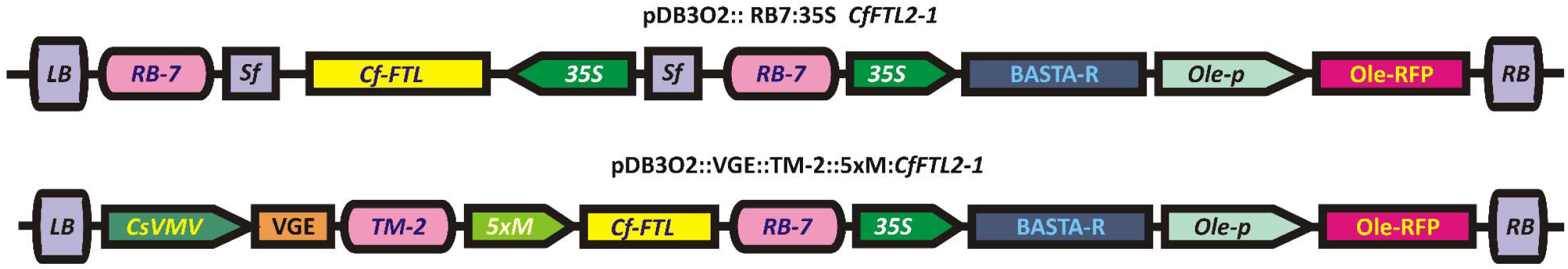
Schematic representation of T-DNA constructs used for the transformation of Arabidopsis. LB, RB – left and right T-DNA borders respectively; RB-7, TM-2 –matrix attachment regions from tobacco; Sf-short stuffer fragment 35 bp, *Cf-FTL* – *C. ficifolium FTL* ORF,, *35S* – Cauliflower mosaic virus 35S promotor; BASTA-R phosphinothricin N-acetyltransferase gene conferring tolerance to Basta herbicide; *Ole-P* - oleosin promotor from Arabidopsis, *Ole-RFP* – gene for RFP reporter protein fused to Arabidopsis oleosin; *CsVMV* promotor from Cassava vein mosaic virus; *VGE* - chimeric transcription factor VGE reactive to methoxyfenozide, *5xM* – minimal *35S* promoter fused with 5 copies of Gal4 binding domain. Not drawn to scale.

### Arabidopsis transformation

The plasmid vectors with a cassette were transferred into *Agrobacterium tumefaciens* strain EHA105 (Hood *et al*., 1993) using the freeze-thaw method of (An, 1987). Arabidopsis wild types (Landsberg *erecta* L*er* or Columbia-0 Col-0) or *ft* mutants (CS56 *ft-1*, Cs185 *ft-3*) were transformed by the floral dipping method (Clough and Bent, 1998). Primary transformants (T1 generation) were selected by spraying 120 mg l^-1^ BASTA^©^ (Glufosinate-ammonium; Bayer, Germany, 150 g l^-1^) three times at 3 - 7 day-intervals, starting with 7 day-old seedlings grown on soil. T1 plants were self-pollinated to produce T2 generation. T2 seeds homozygous for the insertion were identified based on the higher intensity of red fluorescent using LEICA microscope (DM5000B) with LEICA CTR5000 light source. T3 progeny was obtained by self-pollination from the T2 homozygous lines. The presence of transgenes was verified by PCR amplification with BAR_F and BAR_R primers, and with the primers targeted to the *FTL* genes (**Supplementary Table S1**).

### Plant growth conditions and phenotypic scoring

Arabidopsis seeds were stratified for 2 days at 4 °C and sown on Jiffy-7 tablets (41 mm diameter, Jiffy Products International AS, Norway). At 10 days, seedlings were transplanted individually to new Jiffy-7 tablets. Plants were grown in a cultivation room under long days (16 h : 8 h light : dark) at 20°C. Transgenic and control plants were grown in cultivation chamber E-36L2 (Percival Scientific, Perry, IA,USA) under 12 h : 12 h light : dark, 130 μmol m^-2^ s^-1^ light intensity, and 70% relative humidity 23 °C at day and 22 °C at night until flowering. To measure flowering time, the number of rosette leaves at bolting was counted. We conducted one-way ANOVA, honestly significant differences (HSD) were determined by Tukey test, implemented in IBM SPSS Statistics.

### *CfFTL2-1* induction in transgenic Arabidopsis

Transgenic plants carrying the *CfFTL2-1* gene under the control of methoxyphenozide-inducible transcription factor (*VGE::TM-2::5xM:CfFTL2-1*), which were capable of reproduction to produce the T3 generation (30 individuals of each line), were subjected to induction treatment. Plants were grown in the Percival growth chamber under cultivation conditions as described above. A solution of 65 µM methoxyfenozide (Integro, Corteva) was sprayed on plants three times with three-day intervals between applications, starting at the 6-9 leaf-stage (at the age of 4 weeks). The control plants were not chemically treated. The same experiment was conducted with untransformed Arabidopsis of the same genetic background as transgenic plants. Leaves for RNA extraction were sampled from six randomly selected plants immediately before the application of methoxyfenozide and from the same plants again at bolting, when leaf number was also determined.

### RNA extraction and cDNA preparation

Total RNA was extracted using the Plant RNeasy Mini kit (Qiagen, Valencia, CA, USA). DNA contamination was eliminated by DNase I treatment according to the manufacturer’s protocol (DNA-free, Ambion, Austin, TX, USA). If necessary, the DNase treatment was repeated to remove any traces of genomic DNA. RNA quality and concentration were checked on a 0.9% agarose gel and by NanoDrop (Thermo Fisher Scientific, Vantaa, Finland). Single-strand DNA (cDNA) was synthesized from 1 μg of RNA at 55 °C for 30 min. RNA was heated together with oligo dT primers (500ng) for 5 min at 65 °C, chilled on ice and mixed with Transcriptor buffer (Roche, Diagnostics, Mannheim, Germany), 0.5 μl of Protector RNase Inhibitor (Roche), 2 μl of 10 mM dNTPs and 10 units of Transcriptor Reverse Transcriptase (Roche).

### Quantitative PCR

qPCR was performed on the LightCycler 480 platform (Roche) with LC SYBR Green I Master (Roche) in a final volume of 10 μl with 500 nM of each of the primers (**Supplementary Table S1**). The program was: 10 min of initial denaturation at 95 °C, then 40 cycles for 10 s at 95 °C, 10 s at 60 °C (at 58°C for *AtUBQ10*), followed by 15 s at 72 °C. Stable expression of the reference gene *AtUBQ10* was confirmed previously as described by Libus and Štorchová (2006). The PCR efficiencies were estimated based on serial dilutions of cDNAs and used to calculate relative expression using the formula E_R_ ^CpR^/ E_T_ ^CpT^, where ET /ER represents the PCR efficiencies of the sample and reference, respectively, and CpT /CpR represents the cycle number at the threshold (crossing point).

## RESULTS

### *CfFTL1*, but not *CfFTL2-2*, accelerated flowering in Arabidopsis

Primary transformants of Arabidopsis in Col-0, L*er* and *ft-3* genetic backgrounds carrying *CfFTL1* under the control of the strong constitutive promoter *35S* flowered early, after forming three to five rosette leaves. However, only some of them were able to produce viable seeds and give rise to further generations (**Table 1**). We estimated leaf numbers and measured expression of the *CfFTL1* transgene numbers in independent lines of T2 and T3 generations (**Fig. 2A**). We found significantly lower leaf number in the transformants compared with recipient, which implied accelerated flowering, only in lines expressing the transgene. Two Col-0 and three L*er* transgenic lines did not express *CfFTL1* and their flowering time did not differ from the respective wild types. Interestingly, two lines flowered earlier than wild type only in the T2, not in the T3 generation, which also correlated with transgene expression in the respective generation. The decline of transgene expression suggests its gradual silencing.

**Table 1.** The numbers of all primary transgenic lines and the lines capable reproduction, obtained by the transfer of the *FTL* genes of *C. ficifolium* and *C. quinoa* to Arabidopsis wild types and *ft* mutants.

| Transgene cassette | Arabidopsis genetic background | Primary transgenic lines | Lines producing seed | Lines producing progeny |
| --- | --- | --- | --- | --- |
| *35S::CfFTL1* | Col-0 | 16 | 12 | 5 |
|  | Ler | 12 | 6 | 3 |
|  | CS185 *ft-3* | 10 | 6 | 2 |
| *35S::CfFTL2-2* | Col-0 | 10 | 10 | 10 |
|  | Ler | 20 | 20 | 20 |
| *35S::CfFTL2-1* | Col-0 | 0 | 0 | 0 |
|  | Ler | 0 | 0 | 0 |
|  | CS185 *ft-3* | 0 | 0 | 0 |
| *VGE::TM-2::5xM:CfFTL2-1* | Col-0 | >100 | 20 | 6 |
|  | Ler | 24 | 4 | 2 |
|  | CS185 *ft-3* | 3 | 0 | 0 |
|  | CS56 *ft-1* | 28 | 3 | 2 |
| *VGE::TM-2::5xM:CqFTL2-1* | Col-0 | >100 | 23 | 15 |
|  | Ler | >50 | 22 | 18 |
|  | CS185 *ft-3* | 3 | 0 | 0 |
|  | CS56 *ft-1* | >50 | 10 | 6 |

**Figure 2.**
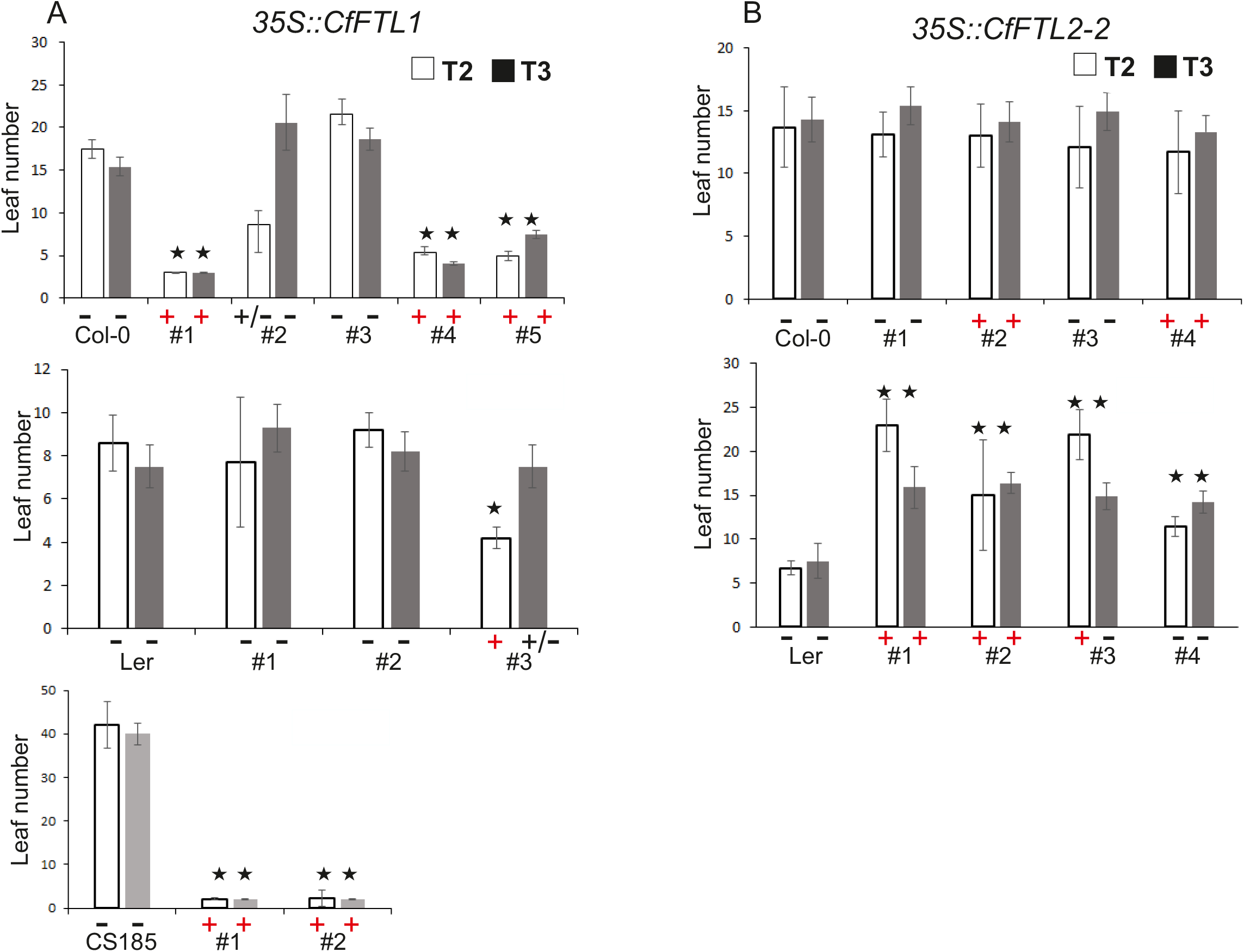
The number of rosette leaves at flowering time in Arabidopsis transformed with the CfFTL genes under 35S promoter in the T2 and T3 generations. A. The CfFTL1 transformants in the Col-0, Ler and CS185 (ft-3) backgrounds. B. The CfFTL2-2 transformants in the Col-0 and Ler backgrounds. The averages and standard deviations were calculated from 20 to 35 plants of the respective independent lineages, which are labeled by the numbers on the x axis. Asterisks represent honestly significant difference (HSD) estimated by Tukey test. The “+” and “– “ signs under the bars indicate the presence or absence of transgene expression (at least five plants analyzed by RT qPCR, with two technical replicates). The “+/-” sign was used, when transgene expression was detected in only one of the individuals of the lineage.

The promotion of flowering was particularly prominent in the CS185 *ft-3* mutant with the *CfFTL1* transgene (**Fig. 2A**). Whereas mutant plants flowered very late after producing about 40 rosette leaves, the transgenic lines flowered early with 4 - 5 rosette leaves, similarly to transgenic plants derived from wild type genetic backgrounds and overexpressing the transgene.

The Col-0 transgenic lines with *CfFTL2-2* under the control of the *35S* promoter flowered at the same time as the Col-0 wild type, while the L*er* transgenic lines flowered slightly later than the L*er* wild type (**Fig. 2B**). The slight delay of flowering in transgenic L*er* plants did not depend on *CfFTL2-2* expression. It could not have been due to the specific activity of this gene, since it harbors a large deletion in its functionally important fourth exon (Štorchová *et al*., 2019).

### Some Arabidopsis seedlings with the inducible *CfFTL2-1* transgene flowered immediately after germination even without induction

We were unable to recover Arabidopsis transformants with the *35S*::*CfFTL2-1* construct, despite repeated floral dipping experiments. Then we noticed several seedlings dying somewhat later after the Basta application. The amplification of their DNA with specific primers (**Supplementary Table S1)** confirmed the presence of the *CfFTL2-1* transgene. Thus, transformation of Arabidopsis with *CfFTL2-1* gene under the *35S* promoter was lethal.

To understand the impact of *CfFTL2-1* on Arabidopsis, we placed this gene to the VGE inducible system (Semenyuk *et al*., 2010), which enables induction of the transgene by methoxyfenozide. Two types of primary transformants were obtained after the transformation with *VGE::TM-2::5xM:CqFTL2-1* (**Table 1**). Some seedlings flowered immediately after expanding cotyledons. They produced tiny flowers, sometimes with well developed stigmas, or small flower buds with prominent trichomes (**Fig. 3**). All these plants died early without producing seed. As no methoxyphenozide was used, premature flowering was most likely caused by the leakage in the *CfFTL2-1* expression. Other *CfFTL2-1* primary transformants did not differ from recipient plants in their flowering phenotype.

**Figure 3.**
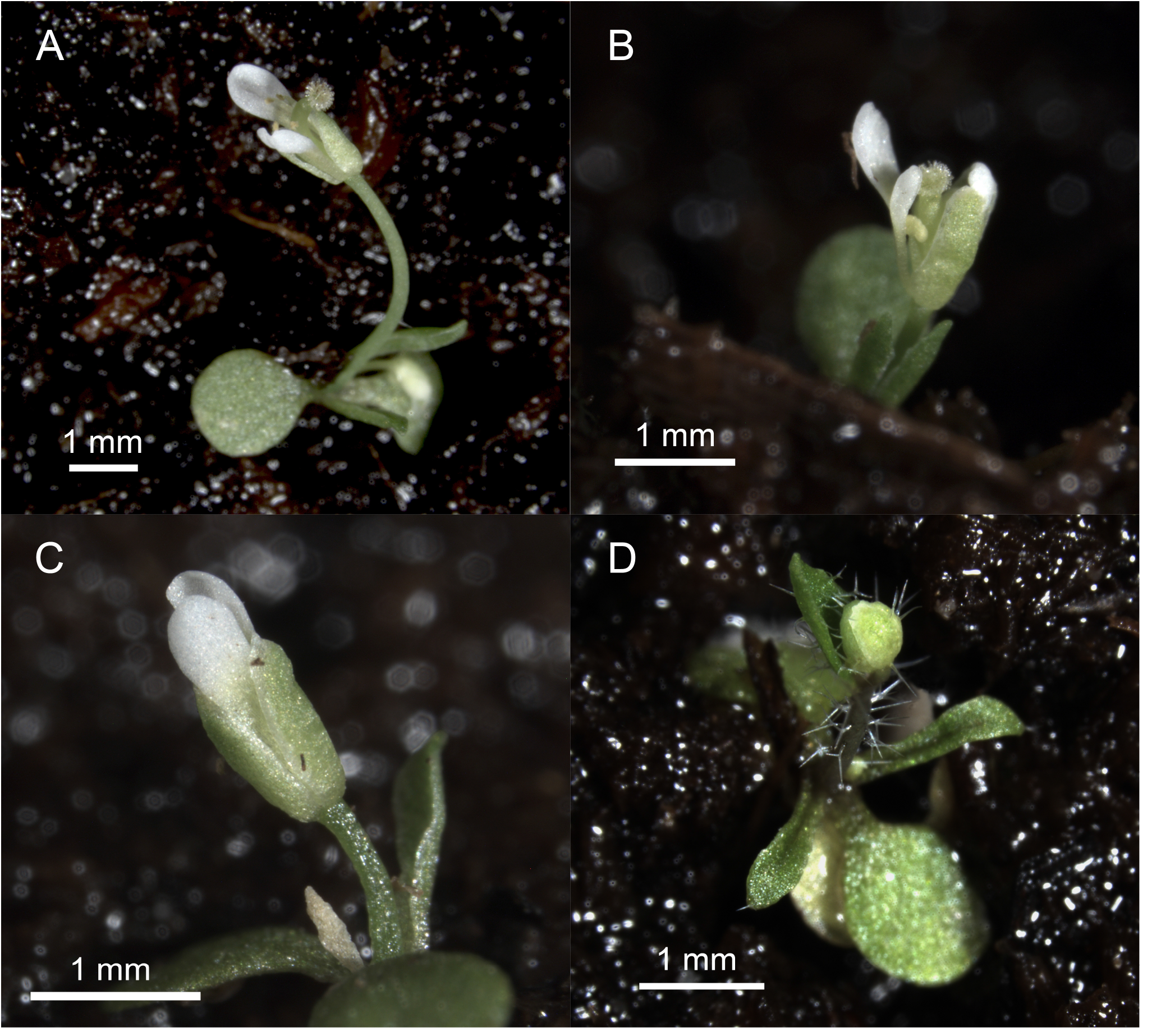
Phenotypes of primary transformants of Arabidopsis Col-0 carrying CfFTL2-1 under the complex metoxyfenozide-inducible promoter (VGE::TM-2::5xM:CfFTL2-1), which flowered without chemical induction. Plants started to bolt immediately after germination. Some of them formed minuscule flowers (A, B, C), others produced tiny flower buds with long trichomes (D). All the plantlets died without generating viable seed. Photo: Lukáš Synek.

The transformation of Arabidopsis with the *VGE::TM-2::5xM:CqFTL2-1* construct, bearing the *FTL2-1* gene of quinoa, provided the same results as transformation with its *C. ficifolium* ortholog. Many primary transformants flowered just after germination and died (**Table 1)**. Some plants gave rise to transgenic lines, in which the transgene was not expressed without methoxyphenozid induction.

### Arabidopsis carrying the inducible *CfFTL2-1* transgene accelerated flowering after methoxyfenozide application

We selected transgenic lines with inducible *CfFTL2-1* in the Col-0 and mutant CS56 *ft-1* genetic backgrounds to effect gene expression by methoxyfenozide. The recipient plants Col-0 and CS56 *ft-1* served as controls. The transgenic plants flowered significantly earlier after the application of methozyphenozide than control or untreated plants (**Fig. 4A**). The acceleration of flowering was accompanied by activation of the transgene. Whereas *CfFTL2-1* expression was negligible before methoxyfenozide application, it increased 100-to 1000-fold after the treatment. The transgene transcript levels differed considerably among individual plants of the same line, as documented by very high standard errors (**Fig. 4B**). However, the plants with both higher and lower *CfFTL2-1* expression flowered approximately at the same time after forming similar numbers of leaves. The rather uniform effect of various transgene transcript levels may be explained by the existence of the threshold value necessary for floral induction. After crossing the threshold, additional *CfFTL2-1* expression did not further accelerate flowering.

**Figure 4.**
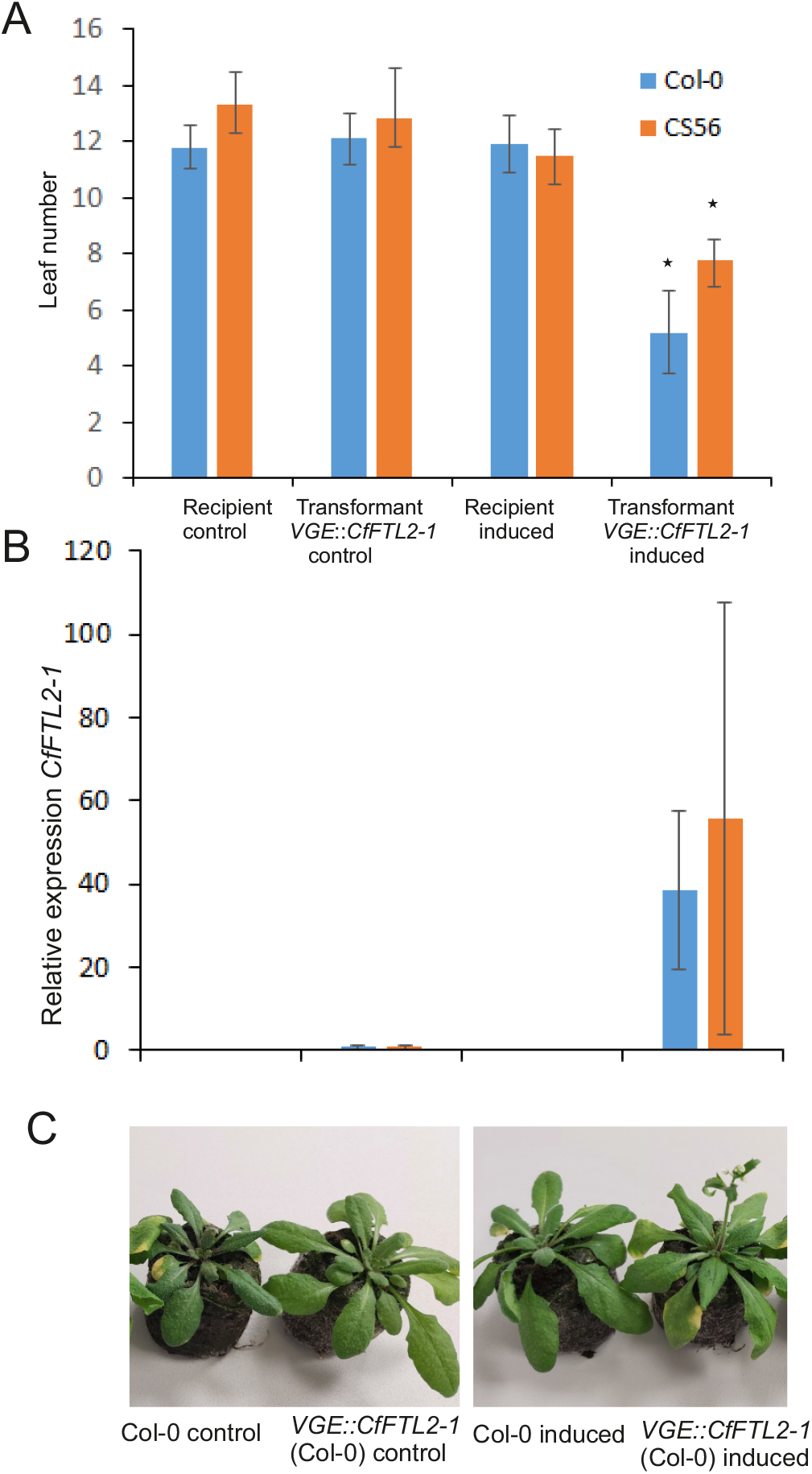
The acceleration of flowering after the induction by 65 µM metoxyfenozide in Arabidopsis carrying the CfFTL2-1 transgene and control plants. A. The number of rosette leaves formed since the time of metoxyfenozide treatment till flowering in Col-0 and CS56 (Ler ft-1) backgrounds, calculated as the average with standard deviation from 20 -30 plants of the same homozygous transgenic line. Asterisks represent honestly significant difference (HSD) estimated by Tukey test. B. The CfFTL2-1 gene expression relative to the reference AtUBQ10 in induced and control plants (the average and standard deviation of 6 individuals) at flowering time. C. The picture of Col-0 and transgenic Arabidopsis plants taken about 9 days after metoxyfenozide treatment.

## Discussion

The *CfFTL1* and *CfFTL2-1* expression in Arabidopsis promoted flowering in both wild types and *ft* mutants, which is consistent with their roles as floral activators. This finding is not unexpected, because the CfFTL1 and CfFTl2-1 proteins share the same sequence with most angiosperm FT floral activators, including the sugar beet floral promoter BvFTL2, in the functionally important region in external loop of the protein (Štorchová *et al*., 2019). They do not possess amino acids Asn(N)134, Gln(Q)141, and Gln(Q)142 responsible for the suppression of flowering in the sugar beet floral inhibitor BvFT1 (Pin *et al*., 2010). The three amino acid substitutions that converted BvFT1 function in sugar beet from the activation of flowering to its opposite most likely occurred after the *Beta* ancestor had diverged from the *Chenopodium* ancestor.

The impact of the *CfFTL1* and *CfFTL2-1* expression on Arabidopsis development differed substantially between the two genes. The overexpression of *CfFTL1*, driven by the constitutive *35S* promoter, accelerated flowering in transgenic lines. In contrast, the overexpression of *CfFTL2-1* was lethal for Arabidopsis seedlings. To estimate its function, we had to place this gene to the inducible cassette and to induce it with methoxyfenozide. Because the selection of primary transformants occurred in the absence of the chemical inducer, we expected the same flowering behavior in both transformants and recipient plants. Surprisingly, a large proportion of transformants started to flower immediately after expanding cotyledons and died without forming seeds. The VGE casette with the *CfFTL2-1* gene is protected against transcription of recipient DNA by tobacco MAR elements (Zhang *et al*., 2002; Allen *et al*., 1996). However, even such isolation from the genomic background is not absolute and may lead to leaky transgene expression in some primary transformants. Premature flowering exhausts resources, which may prevent transgenic plants from the production of viable seeds. The sudden reprogramming from vegetative growth to the reproduction in the very early developmental stage can be also responsible for the lethality of the *CfFTl2-1* overexpression driven by the *35S* promoter. In this case, seedlings died just after germination, having formed only cotyledons.

Transformation of Arabidopsis with the *CfFTL1* gene under the control of the *35S* promoter produced some permanent transgenic lines, but many primary transformants did not produce viable seed. The successfully reproducing lines often silenced the *CfFTL1* transgene. It is therefore possible that the *CfFTL1* overexpression also reduced reproduction owing to premature induction of flowering similarly to the *CfFTL2-1* transgene, while its effect on promoting vegetative growth was much weaker.

The *CfFTL2-1* gene is one of two products of the *FTL2* gene duplication, which occurred after the divergence of *Chenopodium* from *Beta*. The second duplicate *CfFTL2-2* harbors a large deletion, which removed 130 bp of the fourth exon including the motifs necessary for the function of this gene (Štorchová *et al*., 2019). Thus, it is not surprising, that the *CfFTL2-2* overexpression did not affect flowering in Arabidopsis. The slight delay in flowering observed in the L*er* background, but not in Col-0, was most likely caused by the insertion of the construct, not by the transgene expression itself. Thus, unlike *CfFTL2-1*, its *CfFTL2-2* paralog does not seem to play any role in floral transition in *C. ficifolium*, most likely due to the large deletion removing functionally important amino acids. However, it is possible that it participates in the regulation of flowering in other *Chenopodium* species, *e. g*. in *C. suecicum*, where it is present in a complete form and is rhythmically expressed during floral induction (Štorchová *et al*., 2019).

The *FT* gene sequences are highly conserved among angiosperms and thus are expected to maintain their function when transferred to phylogenetically unrelated species. For example, overexpression of *PnFT1* of *Pharbitis nil* (Hayama *et al*., 2007), *CrFTL1* of *O. rubra* (Cháb *et al*., 2008), *BvFT2* of sugar beet (Pin *et al*., 2010), or *GmFT2a* of soybean (Sun *et al*., 2011) promoted flowering in Arabidopsis. However, we have not found any report of lethality caused by the ectopic expression of an angiosperm *FT* gene in Arabidopsis. The underlying cause of lethality due to *CfFTL2-1* overexpression may be immediate floral induction during germination of Arabidopsis seedlings, which is stronger and faster than the activation of flowering controlled by other angiosperm *FT*s, including *CfFTL1*. Because *Chenopodium* is recalcitrant to stable transformation with *Agrobacterium*, we were unable to effect transformation in a homologous system. We are currently running the experiments with virus-induced gene silencing in *Chenopodium* to confirm our conclusions.

If *CfFTL2-1* does act as a powerful promoter of flowering, then we may better understand the results of the study of photoperiodic floral induction in *C. ficifolium* (Štorchová *et al*., 2019). The accession 459 highly upregulated *CfFTL1* under short days, when its flowering was accelerated, too, which was consistent with the promotional role of this gene. In contrast, the long-day accession 283 flowered earlier under long days without apparent activation of any *FTL* gene. However, when *CfFTL2-1* encodes a very strong promoter of flowering, even a very low increase in *CfFTL2-1* transcription, not detected by RT qPCR, could accelerate flowering under long days. We are now testing this hypothesis by the comprehensive analysis of the global transcriptomes during photoperiodic floral induction in *C. ficifolium* 283.

*Chenopodium ficifolium* was proposed as a potential diploid model species for the genetic analyses of the tetraploid crop quinoa (Subedi *et al*., 2021). We have therefore transformed Arabidopsis with the inducible *CqFTL2-1* gene of quinoa to see whether this would result in the same outcome as the transfer of its *C. ficifolium* ortholog. The CfFTL2-1 and CqFTL2-1 proteins differ only in two amino acid substitutions, located outside the functionally important regions.

Hence it is not unexpected that the transformation of Arabidopsis with inducible *CqFTL2-1* would produce the same results as with the inducible *CfFTL2-1* gene; namely, the appearance of many tiny, precociously flowering primary transformants. This experiment supports the usefulness of *C. ficifolium* as a model to be compared with quinoa. The identification of floral activators of quinoa may also have practical importance for quinoa breeding, particularly as the crop spreads to areas of the globe where short- and long-day flowering responses would be advantageous for increasing yields through heat-stress avoidance during the normal flowering period.

Our results are interesting from the perspective of the evolution of gene function. The BvFT1 protein became the repressor of flowering after the genus *Beta* had diverged from *Chenopodium* within the Chenopodiaceae-Amaranthaceae, whereas the CfFTL2-1 protein retained its original function as a floral promoter. The prevailing hypothesis is that activation of flowering was the most likely ancestral role of FTL2 because CfFTL2-1 shares three functionally important amino acids with FT activators, not with its ortholog BvFT1.

The *FTL2-1* genes of *C. ficifolium* and *C. quinoa* triggered precocious flowering in Arabidopsis seedlings despite being homologs of the *BvFT1* floral repressor. This finding illustrates the distict evolutionary trends of two *FT* paralogs which diverged early during the evolution of in the family Amaranthaceae (Drabešová *et al*. 2016). The *BvFT2, CfFTL1* genes and their orthologs retained a conserved gene structure and floral activator function. In contrast, *BvFT1, CfFTL2-1, CfFTL2-2* and their orthologs underwent prominent structural changes including exon acquisition, large deletions or complete loss, and functional diversifications. Thus, the *FT1*/*FTL2* lineage became a versatile toolkit of the evolution enabling the adaptation of annual fast-cycling amaranths to variable environments.

## Supplementary data

The following supplementary data are available at JXB online.

*Table S1*. Primers used in qPCR and for amplification and domestication of *FTL* genes

*Table S2*. DNA components used for the construction of plasmids for permanent transformation of Arabidopsis

## Acknowledgements

The authors are grateful to Lukáš Synek for the photo of tiny flowering Arabidopsis plants. They acknowledge the valuable comments and linguistic help by Eric N. Jellen.

## Author contributions

HS: conceptualization; OAJA, CB, TM, D G-L, KE, ZV and HS: methodology; OAJA, MK and DG-L: resources; HS: writing - original draft; OAJA, DG-L, TM, MK and HS: writing - review & editing; OAJA and HS: supervision; HS: funding acquisition.

## Data availability

GeneBank accession numbers of all the genes are given in the text. The seed of transformed lineages are available upon request.

## Abbreviations

CaMV: Cauliflower Mosaic Virus
*FT*: *FLOWERING LOCUS T*
*FTL*: *FLOWERING LOCUS T like*
MAR: Matrix attachment region
*UBQ*: *UBIQUITINE*

